# Nationwide Inclusive Facilitator Training: Mindsets, Practices and Growth

**DOI:** 10.1101/2024.01.08.574770

**Authors:** Diane Codding, Alexandria H. Yen, Haley Lewis, Vanessa Johnson-Ojeda, Regina F. Frey, Sarah Chobot Hokanson, Bennett B. Goldberg

## Abstract

Advancing diversity in STEM requires competent and confident faculty and staff who can lead local professional development in inclusive teaching to improve classroom instruction and support all learners. This paper examines how a facilitator training model designed to promote inclusive facilitation impacted inclusive learning community facilitator self-reported confidence and practices. This mixed methods study reports on survey data from project trained facilitators (*n*=71) collected over four course runs. Facilitators reported significant increases in confidence, with the largest effect sizes in areas related to diversity, equity, and inclusion (DEI) and identity. Qualitative findings indicate the training model effectively aligned facilitators with our approach to inclusive facilitation. Findings demonstrate that professional development in inclusive teaching, and by extension in other equity and diversity topics, can be successfully done at a national scale by centering identity, power, and positionality while upholding ‘do no harm.’ This paper provides a strategy for how DEI-focused faculty development efforts can select, train, and support facilitators on a national scale while maintaining high fidelity to project values and goals.

---

Inclusive teaching requires more than good intentions; it is an ongoing commitment to learning, reflecting, and implementing equitable and inclusive pedagogical practices to support all students. Equitable teaching practices increase students’ sense of belonging (AIP/TEAM-UP, 2020), motivation and engagement (Fink et al., 2018), and self-association with a positive identity in science, technology, engineering, and math (STEM) (Zumbrunn et al., 2014). Faculty have been identified as critical leaders in creating inclusive climates in STEM classrooms (Canning et al., 2019; Handelsman et al., 2022), yet evidence suggests inclusive teaching professional development reaches a select few (Addy et al., 2021; Dewsbury, 2017).

The Inclusive STEM Teaching Project (ISTP) has disseminated a large-scale, open online course through edX that centers power, privilege, and identity to advance the awareness, self-efficacy, and ability of STEM faculty, postdocs, graduate students, and staff to cultivate inclusive learning environments (Calkins et al., 2024). The online, asynchronous course is accompanied by synchronous, course-associated learning communities (LCs) supported by project-trained facilitators, resources, and activities. LC participants engaged in facilitated discussions to advance self-reflection, skill building, and implementation of inclusive teaching practices. LC facilitators received support through an extensive infrastructure developed by the ISTP that aligned project core principles and pedagogies, helped facilitators address challenges, and shared approaches across dozens of simultaneous LCs running nationwide.

We explore how our facilitator training program informed LC development and facilitation based on data from LC facilitators (*n*=71) collected from 50 different LCs in various institutional contexts held over four iterations of the ISTP course. Our mixed methods examination addresses the following research question: *How does the ISTP training cycle (i.e., training and facilitation) impact facilitators’ self-reported confidence and practices in facilitating an inclusive teaching LC?* Our findings can be applied to other LC models, including those beyond online courses or focused specifically on inclusive teaching. As we show, a structure like ISTP, which utilizes an intentionally constructed flexible learning platform together with training, community development, and support of a cohort of facilitators, can be very effective at delivering high-fidelity professional development efficiently, locally contextualized, and widely accessible.

## Background and Literature Inclusive

### STEM Teaching Project

Scholarship on improving STEM learning and teaching in higher education has forefronted the need for greater teaching professional development for current and future faculty (Austin, 2010, 2011; Beach et al., 2012). The Inclusive STEM Teaching Project (ISTP) is a professional development initiative designed to engage mostly faculty, as well as staff, postdoctoral scholars, and doctoral students in developing the knowledge, skills, and mindsets necessary for effective and inclusive STEM teaching. The project centers identity, power, privilege, and positionality across differentiated learning spaces to create “productive discomfort” for learning (Bezrukova et al., 2016; Taylor & Baker, 2019) while upholding the principle of ‘do no harm’ (Rhodes et al., 2009; Tajima, 2021). ‘Do no harm’ refers to intentional actions ISTP takes to avoid putting marginalized individuals in situations that may cause them to re-experience pain or ask them to explain their lived experiences to majority identities.

Productive discomfort pushes instructors, especially those of majority identities, to reflect and develop an awareness of how their teaching practices impact students’ experiences and sense of belonging in STEM (AIP/TEAM-UP, 2020; Fink et al., 2018; Handelsman et al., 2022; Zumbrunn et al., 2014). In terms of outcomes, semi-structured interviews (*n*=80) with participants of the ISTP showed positive growth concerning their awareness of and mindsets toward inclusive teaching ideas and practices, confidence to implement inclusive teaching, validation of their beliefs surrounding inclusion, and their application of inclusive teaching in their classrooms (Hill et al., *under review*).

While ISTP uses an online, asynchronous curriculum like other large-scale training initiatives, the project differs in form and focus. The Center for the Integration of Research, Teaching, and Learning (CIRTL) Network’s STEM Teaching Massive Open Online Course (MOOC) also has provided evidence-based pedagogical training to over 14,977 participants (Goldberg et al., 2023a). However, their training on inclusive teaching is limited to a single module. ISTP is most like the courses developed at Columbia (2019) and Cornell (2020), in that we offer an online course that focuses on inclusive teaching and offers a certificate of completion. Unlike the Columbia and Cornell courses, ISTP is distinctive in its use of synchronous LCs run by project-trained facilitators and project-provided resources. By Fall 2023, ISTP had trained 396 facilitators in teams from 123 different institutions who have run 95 LCs with 770 participants over five iterations of the asynchronous online course. Not only has ISTP structured an initiative that aims to fill a professional development gap in STEM education, but it has also disseminated to a broad audience and created a community of leaders in institutions nationwide to continue to sustain efforts in advancing inclusive learning environments in higher education.

### Training Learning Community Facilitators

LCs are effective in introducing new pedagogical practices to higher education faculty (Furco & Moley, 2016; Gehrke & Kezar, 2016; Nadelson et al., 2013; Tinnell et al., 2019). When paired with at-scale, asynchronous online learning, LCs further help create a motivating and participatory learning environment. Typically, LCs are institution-based, attended and led by faculty and staff from the institution, without specific training (Cox, 2004). ISTP adapts this approach for a national scale by recruiting and training locally-based facilitators from institutions across the country who go on to develop and co-lead local LCs. However, properly training and supporting a nation-wide group of facilitators to confidently lead discussions on diversity, equity, and inclusion (DEI) topics raises significant challenges. While research has shown the success of large-scale training models at improving facilitator confidence (Pfund et al., 2009; Pfund et al., 2017; Rogers et al., 2018), they again differ from ISTP in training facilitators to teach curricula focused on mentorship skills, whereas ISTP focuses on identity-based DEI topics that occur in higher education classrooms. These approaches also differ in length and delivery; the *Entering Mentorship* facilitator training occurred over five days, six hours per day, for a total of thirty hours of facilitator training time (Pfund et al., 2009). ISTP facilitator training was delivered virtually over two days for a total of six hours of training. The ISTP originally shifted to synchronous virtual delivery in response to the COVID-19 pandemic and ultimately found this model to be effective and accessible for delivering nationwide facilitator training. This approach challenges the widely held notion that effective DEI training necessitates a lengthy, in-person training model.

### Facilitator Practices in Online Course-Associated Learning Communities

Our work adds to the literature on online, course-associated LCs by exploring the ways in which facilitators implemented training materials, facilitated DEI conversations, and cultivated spaces of productive discomfort to advance equity and inclusion, and, due to the scale, also allows us to examine fidelity of implementation. Research in this area has been limited, with only a small set of recent studies focusing on facilitation approaches and development in online course-associated LCs (Blum-Smith et al., 2021; House et al., 2023; Martin et al., 2022; McDaniels et al., 2016). In a recent study, House et al. (2023) identify best practices for culturally responsive facilitation when leading DEI training for faculty. They recommend engaging in active listening, modeling proper attitudes and behaviors to participants, and encouraging an environment of productive discomfort. In another study, Blum-Smith et al. (2021) described two approaches to facilitation in online course-associated LCs, strategies which they describe as “stepping in” (i.e., active facilitation actions) as opposed to “stepping back” (i.e., passive facilitation, meant to give participants greater agency). Similarly, Martin et al. (2022) identified a shift from facilitator-focused actions (e.g., facilitators sharing experiences or offering solutions) to participant-focused actions (e.g., facilitators summarizing and amplifying participant statements) as the LC developed over time. Based on a mixed-methods study of CIRTL’s mentor training synchronous online LC, a different modality, McDaniels et al. (2016) found that participants felt more valued and included in their LCs when facilitators emphasized the importance of group dynamics, provided various means of participation, and actively found commonalities amongst participants from diverse backgrounds and identities. Generally, inclusive facilitation practices in LCs were typified by multiple means of encouraging participation, creating opportunities for participant leadership and agency, and adapting to participant needs.

### Facilitator Training Model

ISTP uses a high-fidelity training model in which project personnel directly select, train, and support facilitators as they lead local LCs (Figure 1). Prior to leading an ISTP LC, facilitators apply to be accepted to participate in six hours of training where they receive a portfolio of facilitation resources, including early access to the full online course, a facilitator workbook, followed by ongoing support from the ISTP team. Facilitators (outside of ISTP team members) were not incentivized or compensated. Given our scale and number of facilitators, compensation wasn’t feasible. Facilitators significant commitment and effort in the absence of compensation or formal recognition is worth noting.

**Figure 1.**
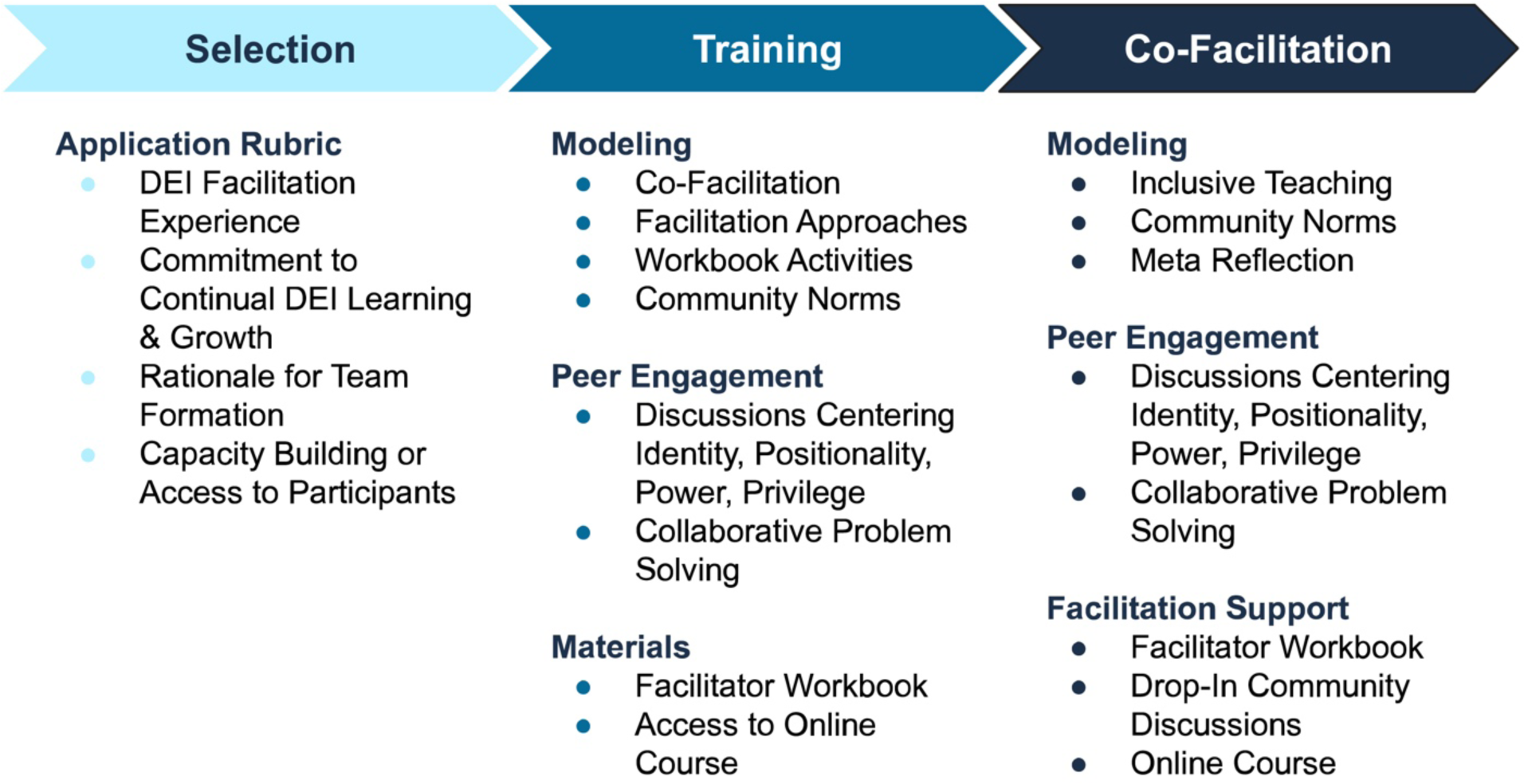
Model for Facilitator Training and Support.

### Facilitator Application and Selection Process

The ISTP training model utilizes team facilitation as a means for encouraging institutions to develop shared capacity for engaging in DEI, as well as to create a local support network (Ortquist-Ahrens & Torosyan, 2009; Wright, 2003). To apply, interested facilitation teams of two to three people submit a combined application that includes a cover letter describing their facilitator team and their interests in the ISTP program, CVs, and individual DEI statements based on the following prompt: *Reflect on why you value diversity, equity, inclusion in your professional and personal life. How do you express your commitment to these values?*

Facilitation teams are evaluated by ISTP personnel on a rubric designed to assess DEI experience and facilitation prior to training, commitment to continued DEI learning and growth, rationale for team formation, capacity building and team’s likely access to participants. These criteria create a common baseline of knowledge, experience, and skill which we believe necessary for upholding the core principle of ‘do no harm.’ Initially, acceptance rates averaged 75%, but have recently increased, indicating that we are reaching our intended audience, with nearly all applicants meeting our criteria of existing experience in and commitment to DEI efforts.

### Facilitator Training

The ISTP facilitator training focuses on identity, power, and privilege within facilitation teams, LCs, and local teaching contexts. Accepted facilitation teams participate in a six-hour synchronous virtual training over two days focused on DEI co-facilitation skills and representative course activities. Grounded in social justice and DEI concepts (Arao & Clemens, 2013; Gillispie, 2018; Goodman et al., 2004; Indigenous Media Action, 2014; Truesdell et al., 2018), our training orientates facilitators to project goals and develops a supportive community. Facilitators experience a subset of our novel course content as participants and then develop their own plans for co-facilitating the activities. In training we model inclusive approaches, such as how facilitators can increase learner agency by guiding rather than leading discussions (Freeman et al., 2014; Lazonder & Harmsen, 2016) and implementing techniques for inclusive and multipartial facilitation (Giacomini & Schrage, 2009; Goldberg et al., 2023b; Routenberg et al., 2013; Zappella, 2007). Structured time is provided for co-facilitators to explore the logistics of running an ISTP LC and facilitation materials (i.e., Facilitator Workbook). Institutional teams train together and are guided to explore local challenges, build collaborative partnerships, and contextualize facilitation for their local setting. Further, three synchronous drop-in community discussions are held virtually during each course run to engage facilitators in reflection activities and crowd-source solutions to current challenges, which are attended by roughly 25% of active facilitators.

### Facilitator Workbook

The ISTP Facilitator Workbook was co-developed by ISTP project team members to provide scaffolded support for teams as they collaboratively plan and facilitate their local LC (Bohrer, 2023). The first sections frame the purpose of LCs and define the roles and responsibilities of LC facilitators, including details on self-reflection, co-facilitation, and collaborative review. The workbook is divided into six modules that were developed to progress in parallel with the online course materials: (1) course overview; (2) diversity, equity, and inclusion in higher education; (3) instructor identity; (4) student identity; (5) inclusive course design; (6) climate in the STEM classroom. Each module includes a summary of the asynchronous course content, LC goals and key takeaways, an introductory activity, two to three central activities associated with the weekly learning goals, and a closing activity. Each activity includes a detailed description and guidance for facilitation. Activities also include prompts, framing questions, and suggested adaptations for different learning contexts (e.g., small or large groups). Each module ends with a debriefing guide.

## Methods

### Data Collection

This study underwent expedited review and was approved by the Northwestern University Institutional Review Board (approval no. STU00207792). Surveys were distributed via Qualtrics to all active facilitators following each course run. The survey consisted of 48 questions with a mix of Likert scale, multiple choice, and open-ended questions (see Codding et al., 2024 for raw dataset and full survey). Questions addressed topics pertaining to facilitation methods and pedagogy, perceived participant experiences, similarity and difference to general DEI facilitation, and utilization of various facilitation resources. The survey explored multiple confidence scales using a retrospective pre- post- approach (Stake, 2002) to examine confidence before facilitator training, after facilitator training, and after LC facilitation. Open ended questions asked facilitators to elaborate on their Likert scale responses and provide insight into their experiences as a facilitator.

The survey items were generated from a grounded approach to examining online course related learning community facilitations (Blum-Smith, et al., 2021), and awareness, confidence, and intent to practice questions (Johnson-Ojeda et al., *under review*). Second, questions were added specifically about the utilization of ISTP facilitator training and resources. Third, expert feedback was provided by ISTP researchers and a small sample of active facilitators. No formal validations, psychometrics or factor analysis was performed.

### Quantitative Data Analysis

Datasets for four course runs were evaluated for this analysis: summer 2021, fall 2021, spring 2022, and fall 2022. All analyses were performed on de-identified data. After the datasets were cleaned in Microsoft Excel, data analysis was run using R version 4.2.2. (R Core Team, 2023), tidyverse (v2.0.0; Wickham, et al., 2019), ggpubr (v0.6.0; Kassambara, 2023a), and rstatix (v0.7.2; Kassambara, 2023b) packages in R version 4.2.2.

For this study, we quantitatively analyzed four survey questions. Three Likert questions retrospectively captured facilitator confidence across three timepoints (pre-training, post-training, and post-facilitation) pertaining to seven areas of facilitation: facilitating DEI conversations, creating open dialogue, creating opportunities for participants to learn from one another, leading conversations centered on identity, leading discussions with higher ed instructors, sharing your own personal narrative, and managing difficult moments in DEI conversations. These questions used a 6-point Likert scale ranging from *Extremely confident* (6) to *Not at all confident* (1). The fourth question asked facilitators to identify how many years they have been involved in DEI-related work using a sliding scale ranging from zero to 25 years.

After evaluating the degree to which the data deviated from parametric assumptions of normality, independence, and outliers (Frost, 2020), we ran paired sample t-tests to determine the growth in confidence of facilitators across time. Cohen’s *d* was used to quantify the practical difference between group means and the relationship between the growth in confidence of facilitators (Cohen, 1969, 1988, 1992). We applied Cohen’s recommendations of *d*≤0.2 small, *d*≤0.5 medium, *d*≥0.8 large for effect sizes (Cohen, 1992). We also compared growth in confidence with years of DEI experience. ANOVA and paired sample t-tests were performed to compare overall group means. A Holm-Bonferroni correction was applied to control the familywise error rate (FWER) in the multiple hypothesis tests and Tukey post-hoc analyses were conducted on the dataset to determine where the differences among the prior DEI years of experience occurred (Wright 1992, 2003).

### Qualitative Data Analysis

We qualitatively analyzed six survey questions that addressed how facilitators created a sense of community, were responsive to LC participants, and encouraged LC participant engagement. We also analyzed questions that asked facilitators to explain what, if anything, they found different in facilitating DEI vs non-DEI-related LCs, how the Facilitator Workbook supported their LCs, and what changes facilitators were planning to make following the LC.

Qualitative analysis was inspired by grounded theory (Glaser & Strauss, 1967), with two researchers independently completing two rounds of open coding and meeting to collaboratively reach a consensus. Emergent codes were organized thematically into parent/child code groups and refined in collaboration with two senior researchers on the project (Strauss & Corbin, 1990). The final thematic codebook included five categories: identity and awareness, inclusive community, LC group dynamics, discussion approaches, and teaching and pedagogy (see Codding et al., 2024). The open-ended survey responses were coded holistically within the context of each survey question, which resulted in the application of a single code unless multiple examples were specified. Qualitative data were analyzed by the first and second authors, both of whom identify as women scholars from majority identities in STEM (white and East Asian, respectively).

Inter-rater reliability (IRR) was conducted on 20% of the qualitative dataset (including non-responses). Inter-rater reliability was calculated Krippendorff’s alpha (*α*) on the ordinal data using the online statistical calculator, ReCal OIR (Freelon, 2013), which accounts for chance agreement. Krippendorff’s (2006) recommends interpreting *α*≥0.80 as indicating robust reliability and *α*≥0.67 as meeting acceptable reliability. In the first round of IRR, *α* values ranged from 0.54 to 1.00. After reviewing questions with an *α*<0.67, we adjusted codes and codebook definitions. A second set of responses representing 20% of the dataset were coded to recheck IRR, with *α* values ranged from 0.64 to 1.00. We reviewed instances where coding did not meet acceptable reliability with *α*<0.67, and then coded the remaining open response data associated with the codebook.

### Participants

We invited all facilitators (*n*=129) who facilitated LCs during our first four course runs (summer 2021, fall 2021, spring 2022, and fall 2022) to participate in this study, 96 of whom completed the survey (response rate 74.4%). We excluded 25 survey participants for either not providing consent or completing less than 50% of the survey. Distinct IDs were assigned to each of the remaining survey respondents (*n*=71).^1^ The cleaned data set included responses from repeat facilitators (*n*=8) who indicated that their most recent facilitation experience was sufficiently different from prior experiences, so each represented a unique data point.

Facilitators applied and were accepted into the program based in part on prior DEI experience. In post-course survey responses, 96% reported attending DEI events, 77% had facilitated DEI events, and 63% had organized DEI events. Figure 2 shows the distribution of the number of years facilitators were involved in prior DEI-related activities, with a mean of 7.37 +/- 5.10 years and a mode of 5.

**Figure 2.**
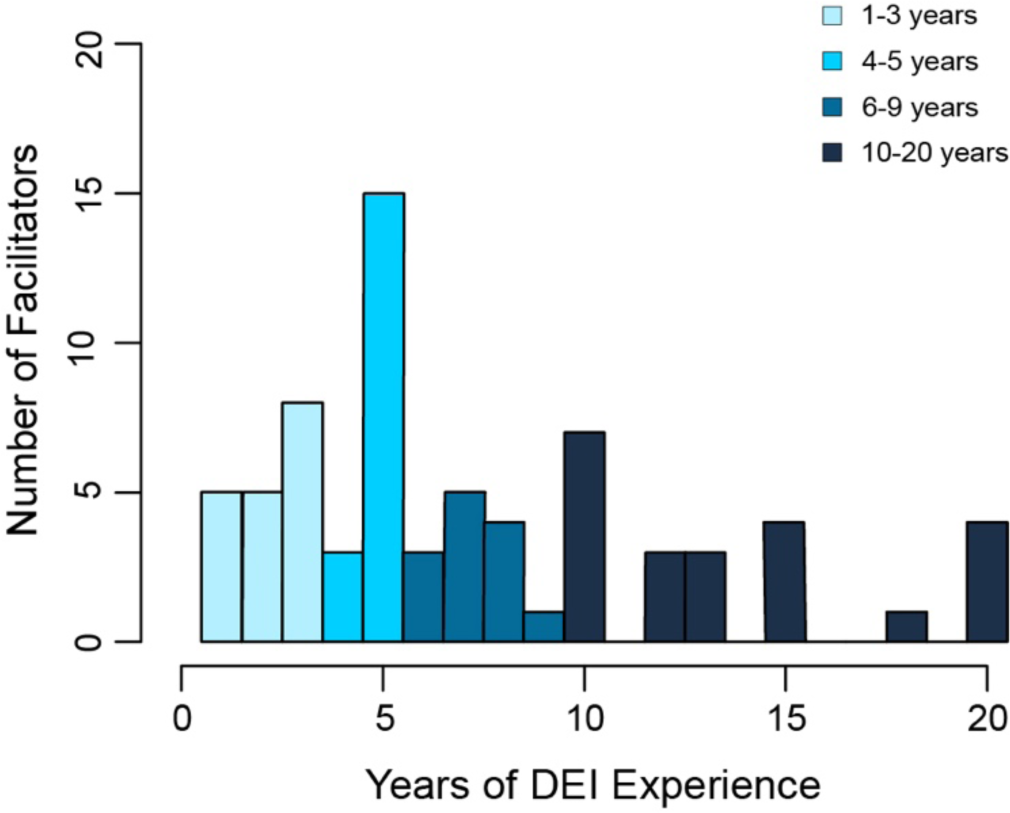
Years Involved in DEI-Related Activities. Note. Facilitators were grouped into quartiles for analysis: 1-3 years (n=18), 4-5 years (n=18), 6-9 years (n=13), 10-20 years (n=22).

## Results

### Retrospective Analysis of Facilitator Confidence

A retrospective analysis shows that confidence increased after facilitators participated in the ISTP training and again after facilitating an ISTP LC (Figure 3). These findings were further supported by paired-sample t-tests with the Holm-Bonferroni correction, which showed that increases in confidence were significant (*p*<0.05) across the seven areas measuring confidence in facilitation. The largest effect size occurred between pre-training and post-facilitation means.

**Figure 3.**
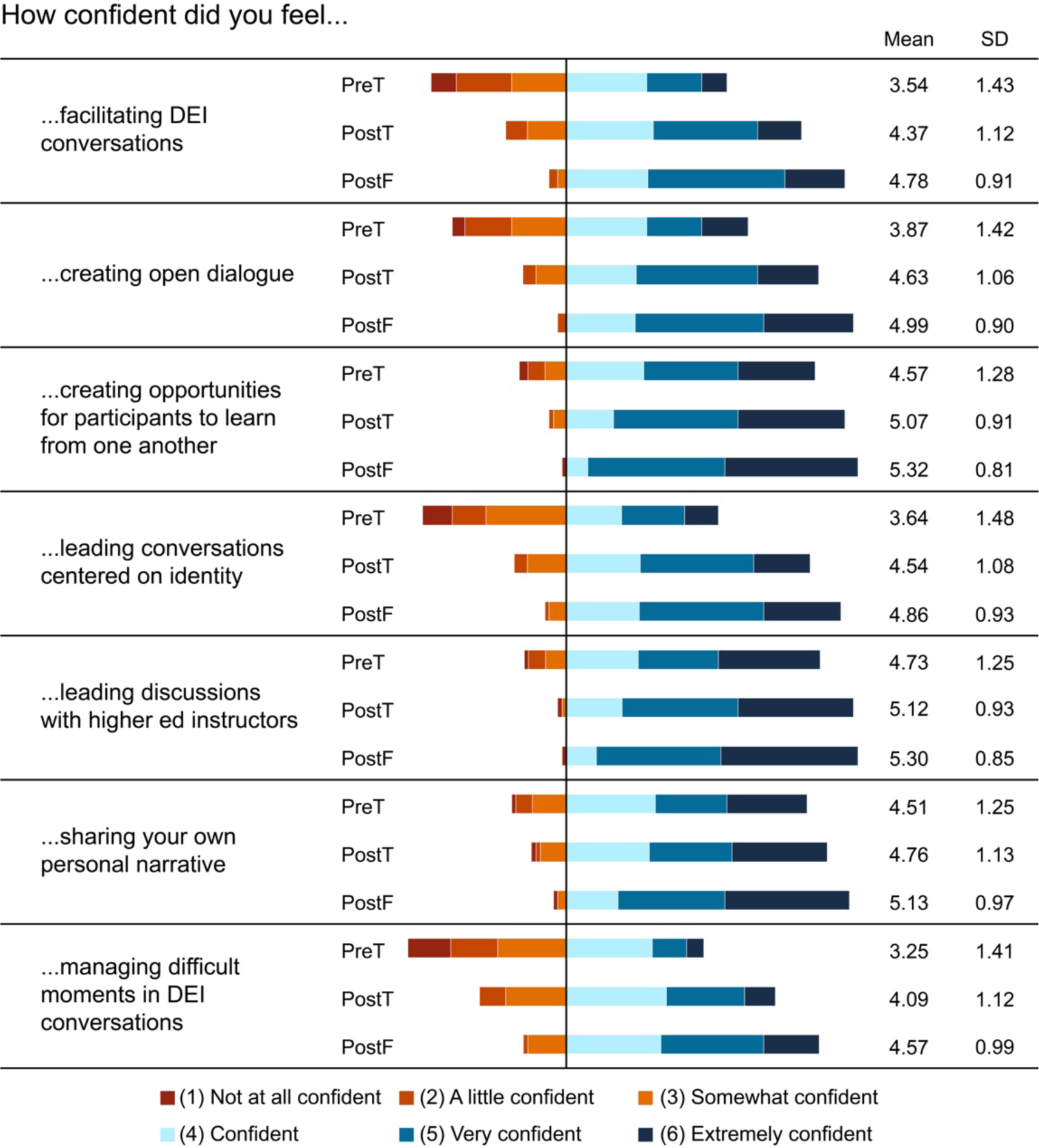
Retrospective Self-Reported Confidence. Note. Data were collected pre-training (PreT), post-training (PostT), and post-facilitation (PostF).

### Change in Facilitator Confidence

As Figure 3 shows, average facilitator confidence was lowest prior to ISTP training, consisting of the highest fraction of “*Not at all confident*” to “*Somewhat confident*” responses. The four areas of lowest average facilitator confidence pre-training were “facilitating DEI conversations,” “creating open dialogue,” “leading conversations centered on identity,” and “managing difficult moments in DEI conversations” (*M*1 = 3.54, *M*2= 3.87, *M*4 = 3.64, *M*7=3.25). The significance of the changes in facilitator confidence were evaluated for all areas, and results showed that the change in confidence for all scenarios (PreT-PostT, PostT-PostF, and PreT-PostF) were significant (*p*<0.05 after Holm-Bonferroni correction) as shown in Table 1.

**Table 1.** Gains of Faculty’s Self-Reported Confidence (paired t-test)

| Question | Time Points (n) | p-value* ($p < 0.05$) | Effect Size | Relative Size |
| --- | --- | --- | --- | --- |
| ...facilitating DEI conversations | PreT-PostT (68) | <0.001 | 0.88 | **Large |
|  | PostT-PostF (68) | <0.001 | 0.66 | Medium |
|  | **PreT-PostF (69)** | **<0.001** | **1.11** | **\*\*Large** |
| ...creating open dialogue | PreT-PostT (68) | <0.001 | 0.83 | **Large |
|  | PostT-PostF (68) | <0.001 | 0.62 | Medium |
|  | **PreT-PostF (69)** | **<0.001** | **1.05** | **\*\*Large** |
| ...creating opportunities for participants to learn from one another | PreT-PostT (68) | <0.001 | 0.62 | Medium |
|  | PostT-PostF (68) | <0.05 | 0.41 | Small |
|  | **PreT-PostF (69)** | **<0.001** | **0.78** | **Medium** |
| ...leading conversations centered on identity | PreT-PostT (68) | <0.001 | 0.90 | **Large |
|  | PostT-PostF (68) | <0.001 | 0.51 | Medium |
|  | **PreT-PostF (69)** | **<0.001** | **1.02** | **\*\*Large** |
| ...leading discussions with higher ed instructors | PreT-PostT (69) | <0.001 | 0.48 | Small |
|  | PostT-PostF (68) | <0.05 | 0.39 | Small |
|  | **PreT-PostF (69)** | **<0.001** | **0.61** | **Medium** |
| ...sharing your own personal narrative | PreT-PostT (68) | <0.05 | 0.37 | Small |
|  | PostT-PostF (68) | <0.001 | 0.57 | Medium |
|  | **PreT-PostF (69)** | **<0.001** | **0.73** | **Medium** |
| ...managing difficult moments in DEI conversations | PreT-PostT (68) | <0.001 | 0.88 | **Large |
|  | PostT-PostF (68) | <0.001 | 0.69 | Medium |
|  | **PreT-PostF (69)** | **<0.001** | **1.16** | **\*\*Large** |

*Note.* **Bolded** cells indicate facilitation scenarios with the largest effect size.
\*p-value adjusted with Holm-Bonferroni correction
Cohen’s $d$ : $d \leq 0.2$ small, $d \leq 0.5$ medium, $**d \geq 0.8$ large

The entire facilitation training cycle (PreT-PostF) saw the greatest effect size in all areas, with “large” relative effect sizes (*d*>0.8) in four areas: “facilitating DEI conversations,” “creating open dialogue,” “leading conversations centered on identity,” and “managing difficult moments in DEI conversations.”

### Change in Confidence Across Years of Prior DEI Experience

Given the effect of training and active facilitation on facilitator confidence, we examined the association of their prior DEI experience with self-reported confidence. We analyzed the changes in confidence over the entire training cycle (PreT-PostF) for each facilitator skill as a function of the number of years of prior DEI experience (1-3 years, 4-5 years, 6-9 years, and 10-20 years) (Figure 4).

**Figure 4.**
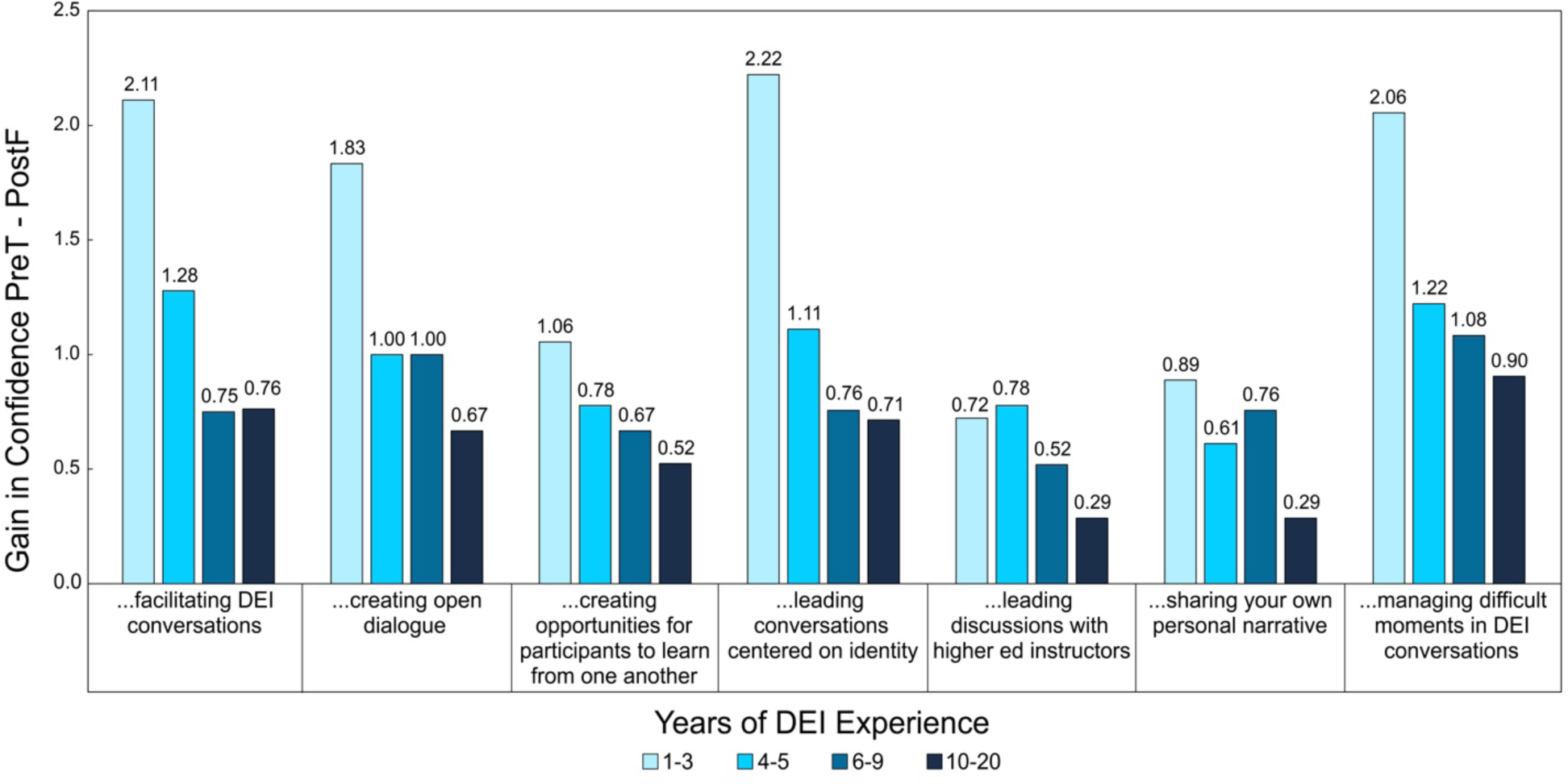
Confidence Across Years of Prior DEI Experience.

The greatest gains in confidence occurred in facilitators who had 1-3 years of DEI experience, with the most growth in the DEI related skills of “facilitating DEI conversations” (gain in confidence, *M*1*: 2.11),* “leading conversations centered on identity” (gain in confidence, *M*4: 2.22), and “managing difficult moments in DEI conversations” (gain in confidence, *M*7: 2.06). Gains in confidence decreased as the years of DEI experience grew, with small differences in gains in most facilitator skills for facilitators with four or more years of DEI experience. The lowest overall gains in confidence occurred in facilitators with 10-20 years of DEI experience, with the lowest gain occurring for “leading discussions with higher ed instructors” and “sharing your own personal narrative” (*M*5 and *M*6: 0.29). All facilitators had small gains in “leading discussions with higher ed instructors,” indicative of their overall experience in higher education. For further analysis, we removed facilitators with 1-3 years of DEI experience to observe if the significant gains had been skewed by their lower average confidence. Using paired t-tests with a Holm-Bonferroni correction, results showed that the change in confidence throughout the entire cycle was still significant (*p*<0.05) but with lower effect sizes (Cohen’s *d* ranged from 0.54 - 0.99). Overall, findings suggested that the combination of ISTP training and actively facilitating an ISTP LC can effectively increase facilitator confidence regardless of their prior DEI-experience. Not surprisingly, facilitators with little prior DEI experience reported the greatest gains in confidence, particularly regarding facilitating DEI conversations.

### Qualitative Analysis of Facilitator Motivation and Reflections

Quantitative results identified *DEI conversations* as a key area of growth in facilitator confidence following the ISTP cycle of facilitator training and LC facilitation. Our qualitative results build on these findings by reporting how facilitators specifically reflected on their own motivations to become ISTP facilitators, and how they used the ISTP materials and training to approach DEI discussions within their LCs. Considering the increase in confidence reported by all facilitators, we also examined facilitators’ self-reported plans to build on their ISTP experiences by pursuing additional DEI activities.

### Facilitator Positionality

ISTP trained facilitators across all years of experience were motivated by a personal commitment to DEI and a desire to contribute to departmental and institutional change. As one facilitator explained, “It is the RIGHT THING TO DO, and I am in a position to make an impact.” Many facilitators felt DEI work was necessary for improving their institutions and advancing faculty pedagogical skills: “We are a minority-serving institution and our students struggle daily with the sorts of experiences defined in the course. My goal is to get every faculty member on campus through this six-week course and set of discussions.” Facilitators also expressed an interest in self-knowledge, hoping to “gain experience,” “expand,” “learn,” and “deepen” their own DEI competency. Facilitators, even those with extensive DEI experience, expressed a desire to improve their existing skills. Experienced facilitators differentiated between their prior DEI experience and our training. One noted that they had previously participated in DEI training sessions but wanted to “become more knowledgeable about these issues [and] become a better facilitator” for the DEI sessions they run. The second facilitator stated, “I wanted to sharpen and broaden my teaching craft with the lens of inclusivity. But I also wanted to learn the skills of facilitation. I wish to become an active listener and also [a] reflective conscientious mediator and strategizer for building a community [for the faculty at my institution].” Findings indicate the ISTP facilitator application process successfully identified facilitators who were intrinsically and extrinsically motivated to engage in DEI work and could specifically describe what skills they hoped to gain from the experience.

### Application of Training and Materials

Facilitator training aligned inclusive facilitation practice through modeling and sharing research-based approaches to the design of the ISTP open online course and pedagogy of the Facilitator Workbook. Most facilitators reported using the Facilitator Workbook during LC facilitation, with 84.3% reporting “moderately” to “extremely” relying on the workbook. Most facilitators, even those with less DEI experience, reported adapting the workbook to suit their participants. As one explained, “We used it as the basis of what we did each week. We made some modifications based on who our faculty were and the time we thought each part would take, but we followed it quite closely.” Overall, facilitators described the workbook as an activity guide that helped to save time and align LC activities with the open online course.

The impact of the ISTP facilitator training was apparent in how closely self-described facilitation practices aligned with ISTP training and were supported by the Facilitator Workbook (Table 1). For example, while facilitators may have had prior experience establishing community guidelines for discussions, findings showed that facilitators intentionally used community guidelines and facilitation practices that emulated the ISTP training. In the first example below (Table 2), the facilitator described introducing guidelines in combination with encouraging participant-led discussions (*ISTP Training*), which removes the facilitator as the expert in the conversation (*Facilitator Workbook*). Additionally, the facilitator added a “real-world application” component by asking participants to consider how they would use inclusive teaching practices in their own classrooms, another frequent component of ISTP training. Several of the ISTP core principles were likely familiar to facilitators prior to training, such as using active and diverse learning approaches, encouraging participants to share in diverse ways, and challenging hesitancies and assumptions. However, findings showed that the *way* in which facilitators applied these skills closely aligned with the resources the project provided and modeled in training, suggesting we acclimated them to specific project-aligned, research-based approaches.

**Table 2.** Alignment of Self-Reported Facilitation Approaches with ISTP Training and Materials.

| Facilitator Response | Facilitator Training | Facilitator Workbook |
| --- | --- | --- |
| In the first week (1) we went over the **community guidelines**, spent some time having the (2) **participants generate their questions and ideas** for what we wanted the guidelines to be in our space, and (3) **how they can generate and invite similar guidelines in their own teaching contexts**. | 1. Community Guidelines<br>2. Participant-Led Discussions<br>3. Planning Real-World Application | 1. The first session sets the tone for participation and sharing space.<br>2. Co-create clear guidelines for how to participate. |
| We used (1) **small group discussion** to allow everyone the chance to engage in topics... (2) **Participants were given the choice to participate in or decline to participate in any topic for any reason**...For some activities, we asked everyone to share, if they were comfortable, in order not to always rely on volunteers for report[ing] out. (3) **We switched up who served as reporter for the small group activities**. | 1. Small Group Discussion<br>2. Acknowledging Diverse Forms of Engagement<br>3. Using Active Learning Strategies | 1. Give participants the option of types of active learning.<br>2. For pair or group work, remind participants that they only have to share what they feel comfortable sharing. |
| [We] solicited input on teaching in their discipline, (1) **asking about language use ("tell us more what you mean when you say unprepared students are a problem for your program")** and (2) **using myself as an example of situations where I changed my mind** from a beginning instructor to now and why. | 1. Addressing Common Hesitancies<br>2. Sharing Experiences | 1. Productively challenge participants to be critical of deficit-minded language; ask for clarification.<br>2. Lean into moments of productive discomfort and offer support. |

*Note.* Facilitators were asked to provide one or two examples of how they regularly encouraged LC participants to engage with inclusive teaching practices.

### Reflections on Leading DEI Discussions and Future DEI Work

To distinguish clearly between facilitating inclusive teaching professional development in our project and non-identity focused teaching professional development, we also asked facilitators to reflect on the differences between leading DEI and non-DEI discussions, and any changes they would implement in a future LC. The consensus among facilitators was that “non-DEI [learning communities] are easier to facilitate than a DEI-related learning community.”

Facilitators emphasized that this was a highly personal, sensitive topic that centered identity and required vulnerable engagement. As one facilitator explained, “I think there was more initial resistance to the topics under discussion than you get with a regular learning community because people came to it with different identities and life experiences. But I think we had more lightbulb moments as a result, too.” Facilitators found that discussions were highly engaging but challenging to facilitate, and participants often hesitated to share.

Facilitators expressed awareness of how their identity, positionality, and privilege affected their LC facilitation. One facilitator reflected on how the act of facilitating an ISTP LC helped them understand the impact of their positionality and privilege:

> I learned that I need to step in more quickly when a microaggression occurs. I made that mistake early on and learned from it. I am also still thinking about the conversation [in ISTP training] around being careful not to offer strategies without being mindful of your own context. I hadn’t realized that this could feel patronizing to some participants until it was brought up. In the future, I will add that to the community discussion guidelines.

Another facilitator noted, “Based on learner feedback, I would preface any suggestion I make as a non-expert suggestion and not the final answer nor applicable to everyone’s situation.” A third facilitator emphasized that in the future they wanted to avoid having facilitators ‘in the lead’ directing conversations.

A core requirement of access to ISTP facilitation resources was team formation; individual facilitators were not accepted. Facilitators identified co-facilitation as a powerful way to mitigate some of the negative impacts of their identity, positionality, and privilege within their LC. As one facilitator explained, “I think it’s really helpful to have a co-facilitator. … For this particular topic, I think it’s really beneficial to have multiple experiences and perspectives available to facilitate the discussions.” Another facilitator elaborated, “We had different skill sets, and it was very helpful. I don’t think I would have felt as comfortable or enjoyed it as much if I was facilitating myself.” Facilitators were keenly aware of both the strengths and challenges that came with the role of facilitator, which were directly connected to activities within the training that were designed to strengthen their team dynamic.

When asked how facilitators would change their future DEI activities after facilitating an ISTP LC, respondents described plans to use ISTP activities and approaches in their classroom and professional development contexts with other faculty. One facilitator reflected:

> I have made progress, but know I need to keep learning. I want to be better at handling difficult conversations and am learning more about that. I also want to try to actively recruit more folks from [my institution] to take this course and participate in a learning community. I’m helping lead some curriculum revision efforts this summer and will be bringing more DEI content to those.

Seven facilitators explicitly stated their intent to facilitate an LC in the future, incorporate ISTP into institutional programming, participate in an LC, or encourage faculty at their institution to take the ISTP course. Facilitators overall reinforced that leading a LC helped them consider more ways to continually develop their understanding of inclusive teaching and DEI.

Leading an ISTP LC seemed to renew their commitment to doing DEI work on an institutional level, especially for facilitators who self-identified as having 10-20 years of DEI experience. As one such facilitator stated, “I am being more forceful in my engagement with the ‘powers-that-be’ at my institution—an institution-wide DEI strategic planning effort is underway and I’m being very pushy about going beyond words on the page to facilitation and monitoring of implementation.” Multiple facilitators stated that they hoped to organize DEI teaching events for their institution and other faculty members. One facilitator shared, “I want to continue for DEI facilitation to be a part of my regular activities. [Facilitating an LC] has reaffirmed my commitment to incorporating students as partners in [the] educational process, for continued attention and improvement in the materials.” While facilitators across all levels of experience affirmed various pedagogical strategies they planned to implement, highly experienced DEI practitioners seemed to feel a particular commitment and enthusiasm for engaging in institution-wide efforts.

### Discussion and Implications

There is an ongoing need in STEM education nationally to engage educators in inclusive teaching professional learning to be able to apply pedagogical practices that broaden student success, particularly among racially minoritized and historically marginalized students (Handelsman et al., 2022). Our goal is to build a nationwide program that trains educators in creating STEM classrooms that retain, support, and motivate diverse student populations. Findings from this study indicate that ISTP was successful in preparing and supporting facilitators to lead LCs using project-aligned inclusive facilitation practices. Our training model significantly increased facilitator confidence related to facilitating DEI conversations, creating open dialogue, leading conversations centered on identity, and managing difficult moments in DEI conversations—four challenges specific to facilitating DEI-focused LCs. Below we discuss key components of our model for facilitator training that have been central to our successful nationwide dissemination.

### Facilitator Selection and Training for Nationwide Scale

There is ample research on the effectiveness of locally focused equity and inclusion faculty development programs (Macaluso et al., 2020; Rogers et al., 2018; Trejo et al., 2022; Womack et al., 2020). However, these programs cannot reach the same national scale achieved by the ISTP through our globally accessible asynchronous online and national LC inclusive facilitator training, with its research-based pedagogical approaches and course-aligned activities—mechanisms we have found essential for successful scaling. To make nationwide implementation possible, ISTP leveraged facilitators’ existing skillsets. We relied on, respected, and valued STEM educators who chose to lead ISTP LCs. By selecting motivated facilitators with an average of seven years of prior DEI experience, we were able to implement a succinct training model (six hours over two days) that focused on norming facilitation practices and equipping facilitators with a carefully curated workbook, rather than developing inclusive facilitation skills from the ground up.

We were able to train facilitators from institutions across the United States to implement our evidence-based inclusive teaching principles successfully in their local LCs. Facilitators demonstrated alignment with ISTP by creating spaces of productive discomfort, upholding ‘do no harm,’ and emphasizing discussions that center issues of identity, power, privilege, and positionality. Findings demonstrate we targeted the right teams of facilitators for leading course-aligned LCs at scale, teams often made up of a combination of STEM faculty members, and DEI and teaching and learning center staff. Additionally, our approach to training has effectively developed inclusive facilitation skills and increased confidence in facilitating DEI-focused LCs. It is also important to note that facilitators reported renewed enthusiasm for and commitment to engaging in DEI work following LC facilitation. Ultimately, we believe we have tapped into a national phenomenon—higher education professional faculty and staff who are looking for a structured, supportive, and high-quality platform with effective but not overbearing training or participation requirements. Our facilitators see the ISTP course and LCs as a means to engage more broadly and deeply across their institutions than yet another implicit bias workshop.

### Increasing Facilitator Confidence through Full Cycle of Support

We have found that providing a full cycle of support (i.e., asynchronous course, facilitator training, materials, and continuing support) is essential for increasing facilitator confidence and skills related to facilitating DEI-focused LCs. Training prepares facilitators to lead LCs, but according to our facilitators, it is the act of facilitating itself that solidifies their inclusive facilitation skills. Facilitating DEI-focused LCs is unique, differentiated by its identity-focus from mentoring and traditional teaching professional development; it requires a specific set of skills for cultivating spaces for productive discomfort and engaging participants in vulnerable discussions around highly personal issues. Evaluating our training process in its entirety revealed that facilitation itself can function as a method of improving DEI confidence. This was particularly true for increasing facilitator confidence related to managing difficult moments during challenging conversations. As one participant explained, “I made that mistake early on and learned from it.” Being able to draw from resources and the expertise within the facilitator community at different time points in our training cycle allowed facilitators to develop themselves and apply their knowledge continuously, as opposed to a single learning moment that had to be extrapolated into practice.

### Limitations

The dataset for this study has three key limitations. First, there was selection bias in the facilitator application process. By only accepting facilitators with some prior DEI experience, less-experienced applicants who fell below our acceptance criteria were excluded. We made this choice intentionally after one training in which we accepted participants with no DEI experience and they self-reported during the training that they were unprepared to lead an ISTP LC. It is possible that with much longer engagement and practice, these novices too could succeed. We also acknowledge that facilitators with an existing foundation of DEI knowledge may be more willing to self-report learning gains as they possess an existing motivation to take the course and develop their DEI skills. Second, data were gathered at a single point, one month after each run rather than across time periods, which limited our findings to a retrospective analysis of confidence gains. Additionally, as data were collected post-facilitation, facilitators who completed the ISTP facilitator training but had not yet facilitated an LC were excluded from the pool of potential study participants. Third, the scope of this paper is limited to facilitators’ self-reported data. However, a recently accepted publication on an individual learning community (Jaimes et al., 2024) has demonstrated significant participant outcomes. Additionally, we are currently in preparation of a manuscript that examines the experiences of participants across more than 40 facilitated LCs with findings revealing a strong alignment between facilitator training, facilitation practices, and participant experiences.

## Conclusion

ISTP was developed to provide professional development for instructors seeking to improve their DEI practices in the classroom at a national scale. To best support these participants and meet the growing demand for inclusive teaching professional development, ISTP utilized a multi-modal learning approach, pairing its open online course with synchronous LCs led by institution-based facilitators. These facilitators, trained by the project and given an extensive Facilitator Workbook, effectively adapted the resources and content provided by ISTP into distinctive LCs addressing local participants’ goals and interests. ISTP has demonstrated that professional development in inclusive teaching, and by extension in other equity and diversity topics, can be successfully done at a national scale by centering identity, power, and positionality while upholding ‘do no harm.’ Further, ISTP has shown that dissemination through project-trained facilitators of local LCs can be successful across a wide range of institutional and disciplinary contexts. This paper provides a strategy for how DEI-focused faculty development efforts can select, train, and support facilitators on a national scale while maintaining high fidelity to project goals.

## Footnotes

1 Data were blinded for data cleaning, analysis, and reporting. After completing our analysis, we learned that our third author had completed the facilitator survey prior to joining the project. This author worked exclusively with quantitative analysis and their qualitative responses were not included as evidence in this paper.

